# Charge and steric hindrance in the glycocalyx govern membrane interactions

**DOI:** 10.64898/2026.06.12.731848

**Authors:** Rafael B. Lira, Jacob Kerssemakers, Cees Dekker

**Affiliations:** Kavli Institute of Nanoscience Delft, Delft University of Technology, Delft, the Netherlands

## Abstract

The outer leaflet of the plasma membrane – the cell’s interface with exogenous biomolecules and particles – is largely electrostatically neutral and highly viscous, conditions that are generally unfavorable for fusion of external membrane-bound objects such as liposomes. Yet living cells efficiently interact and fuse with charged liposomes, revealing a discrepancy between synthetic and biological membranes. Here, we identify the glycocalyx, the carbohydrate-rich layer that coats cells, as the component that explains this mismatch. Using synthetic membranes reconstituted with an artificial glycocalyx, we resolve how electrostatics and steric hindrance regulate the interactions of cationic liposomes with synthetic membranes. Charged glycolipids are found to be sufficient to convert non-fusogenic neutral membranes into fusion-competent membranes. In contrast, large polymers in the glycocalyx suppress fusion. Steric hindrance acts primarily at the docking stage, while fusion proceeds through a long-lived hemifusion intermediate. Strikingly, these large differences occur without measurable changes in membrane viscosity or phase state, demonstrating that glycocalyx-mediated control is independent of the membrane mechanics. Selected glycocalyx components can also promote endocytic budding and cargo internalization. We thus show that the glycocalyx is a tunable physicochemical regulator of liposome docking, fusion, and internalization. The findings provide a framework for understanding how biological agents and therapeutics engage cell membranes and they offer key design principles for synthetic cells.

## Introduction

The glycocalyx, the carbohydrate-rich layer that coats cells, plays a vital role in cell-cell recognition, communication, and protection. It is composed of a lipid (fatty acid) portion that is embedded in the cell membrane and a carbohydrate (sugar) portion that extends out into the extracellular space^1,2^. The glycocalyx is typically composed of a mixture of glycolipids (which are small), glycoproteins (moderate size), polysaccharides (broad size range), and proteoglycans (very large)^3^. It contains a significant proportion of anionic moieties, such as sialic acids and sulfated glycosaminoglycans^4,5^. Under physiological conditions, the glycocalyx is negatively charged. This ensures that the cell surface remains negative, even in the absence of negatively charged phospholipids in the outer leaflet. The glycocalyx has been shown to be involved in the internalization of several natural and synthetic nanoparticles via specific and non-specific interactions^6,7^, and to function as electrostatic receptors for cationic nanoparticles, while at the same time, shielding interactions with anionic ones^8^. Stripping off glycocalyx components from the membrane significantly decreased the uptake of cationic^8^ but not of anionic nanoparticles^6,9^, without interfering with cellular endocytic machinery^9^. While electrostatic interactions thus appear to serve as a selective filter for interactions with external objects, the thick dense structure of the glycocalyx also acts as a physical barrier, sterically restricting access to the plasma membrane and surface receptors^10^.

The glycocalyx extends out of the plasma membrane (PM). The inner leaflet of the PM is rich in unsaturated and anionic phospholipids, conferring it a fluid character and a net negative charge, whereas the outer leaflet is rich in long, saturated, and neutral phospholipids, yielding a more packed structure that is electrostatically essentially neutral^11^. Because fusion of membrane bound organelles such as exosomes or engineered lipid-based nanoparticles such as liposomes, is favoured into fluid membranes, the inner leaflet is more fusogenic than the outer^12^. Furthermore, for many membrane-based systems, fusion with the PM depends on membrane charge^13,14^. Since the lipid composition of the external membrane leaflet is virtually neutral, other components must contribute to the overall negative charge of the cell surface for electrostatic interactions with liposomes, such as the glycocalyx.

The limited understanding of the interactions between liposomes and the glycocalyx-coated cell surface has precluded efficient intracellular delivery of cargos to cells, such as in drug delivery of medicines and genes^7–9^. Liposomes entrapping drugs have been used in clinical practice^15^, and liposome-gene complexes are used as vaccine vectors^16^. Although particle binding and uptake have been relatively well characterized^17,18^, liposome fusion, either directly at the plasma membrane^19^, or with endosomes upon internalization^20^, is much less understood. Drug delivery mediated by direct liposome fusion with the PM has the potential to readily and quickly deliver cargos directly in the cell cytosol. Unlike the slow (hours to days) and inefficient (≈ 1% cargo) intracellular delivery from internalization-based pathways^21,22^, direct fusion of cationic liposomes with cells has been reported to take place in sub-seconds and to deliver large amounts of cargos in the cytosol^23^. Despite efficient fusion with neutral and viscous PM of living cells, cationic liposomes require anionic membranes to fuse with synthetic membranes^24^, and fusion is further inhibited as membrane viscosity increases^25^. We hypothesize that the glycocalyx, present in cells (but absent in typical experiments on synthetic membranes) may be the missing factor enabling efficient liposome binding and fusion with the plasma membrane of living cells.

To resolve the discrepancy between efficient fusion of liposomes with living cells and their poor fusion with neutral, viscous synthetic membranes, we reconstituted liposome–membrane interactions using an artificial glycocalyx. Model membranes provide a controlled environment to isolate the specific contributions of interfacial components while avoiding cellular complexity. As a minimal glycocalyx mimetic, we incorporated glycolipids of defined molecular weight, charge, and grafting density to independently tune electrostatic interactions and steric hindrance. Charged glycolipids were found to sufficient to convert otherwise non-fusogenic neutral membranes into fusion-competent ones, whereas large polymers strongly inhibit membrane fusion through steric hindrance. Single-particle measurements show that this steric inhibition primarily acts at the docking stage, while also slowing down the fusion kinetics of bound vesicles. In addition, membrane bending enables internalization of membrane-bound particles irrespective of fusion, providing an alternative uptake pathway. Notably, these effects occur without detectable changes in membrane phase state or viscosity.

Previously, we found that cationic liposomes fuse highly efficiently with negatively charged model membranes, but mainly dock (and not fuse) to electrostatically neutral giant unilamellar vesicles (GUVs)^13,24^, highlighting the critical role of membrane charge in overcoming the docking-to-fusion barrier. Fusion efficiency is also inhibited by an increase in membrane viscosity^25,26^. These findings contrast sharply with fusion to living cells, as their outer plasma membrane leaflet is very viscous and contains very low fractions of anionic phospholipids, yet still supports efficient fusion, accompanied by decreased surface tension and pronounced inward tubulation^23^. This suggest a discrepancy between minimal synthetic membranes and living cells.

This study aims to determine whether an artificial glycocalyx mesh at the surface of cell-like membranes facilitates fusion of cationic liposome nanoparticles with otherwise non-fusogenic synthetic membranes, how it modulates fusion intermediates, and whether the membrane mechanics contributes to this process. We estimate that electrostatic factors at the cell surface mediate liposome–cell interactions independently of phospholipid charge. We hypothesize that the glycocalyx provides the missing electrostatic and steric regulation required for efficient liposome fusion. To test this, we studied an artificial glycocalyx mimetic system wherein we systematically varied the glycocalyx composition and quantified its effects on fusion efficiency, intermediates, and membrane remodeling. The data confirm the hypothesis and show that the glycocalyx is a tunable physicochemical regulator of liposome docking, fusion, and internalization.

## Results

We reconstituted an artificial glycocalyx in neutral GUVs made of POPC:cholesterol:glycolipid (50:30), or negative GUVs made of POPC:POPS:Chol:glycolipid (30:20:30:20), and probed liposome binding and fusion at the single GUV level. As model glycolipids, we used a variety of structures: small cerebrosides (CBs) and sulfatides (SFTs), with CBs being neutral and SFTs carrying a single negative charge; PEG350 and PEG5000, each carrying one negative charge but differing largely in polymer size; and GQ1b as intermediate in size and carrying four anionic charges (Figure 1A). Depending on grafting density, long and flexible glycolipid polymers may adopt distinct conformations^27^, which we hypothesize will interfere with liposome–membrane interactions (Figure 1B). The glycolipids molecular weights and charge densities are shown in Figure 1C and Table 1.

**Table 1.**
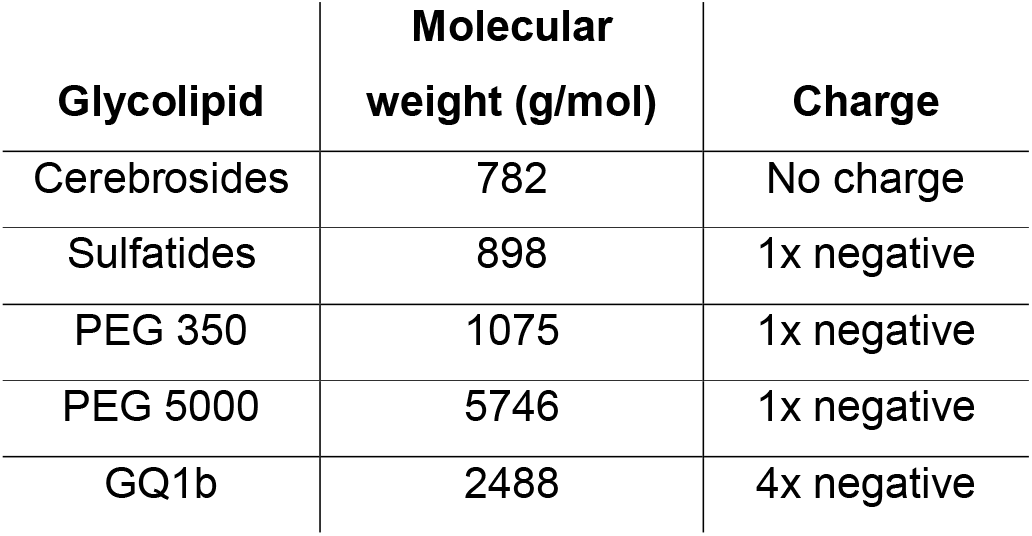
Molecular sizes and charges of the glycolipids used in this study.

**Figure 1.**
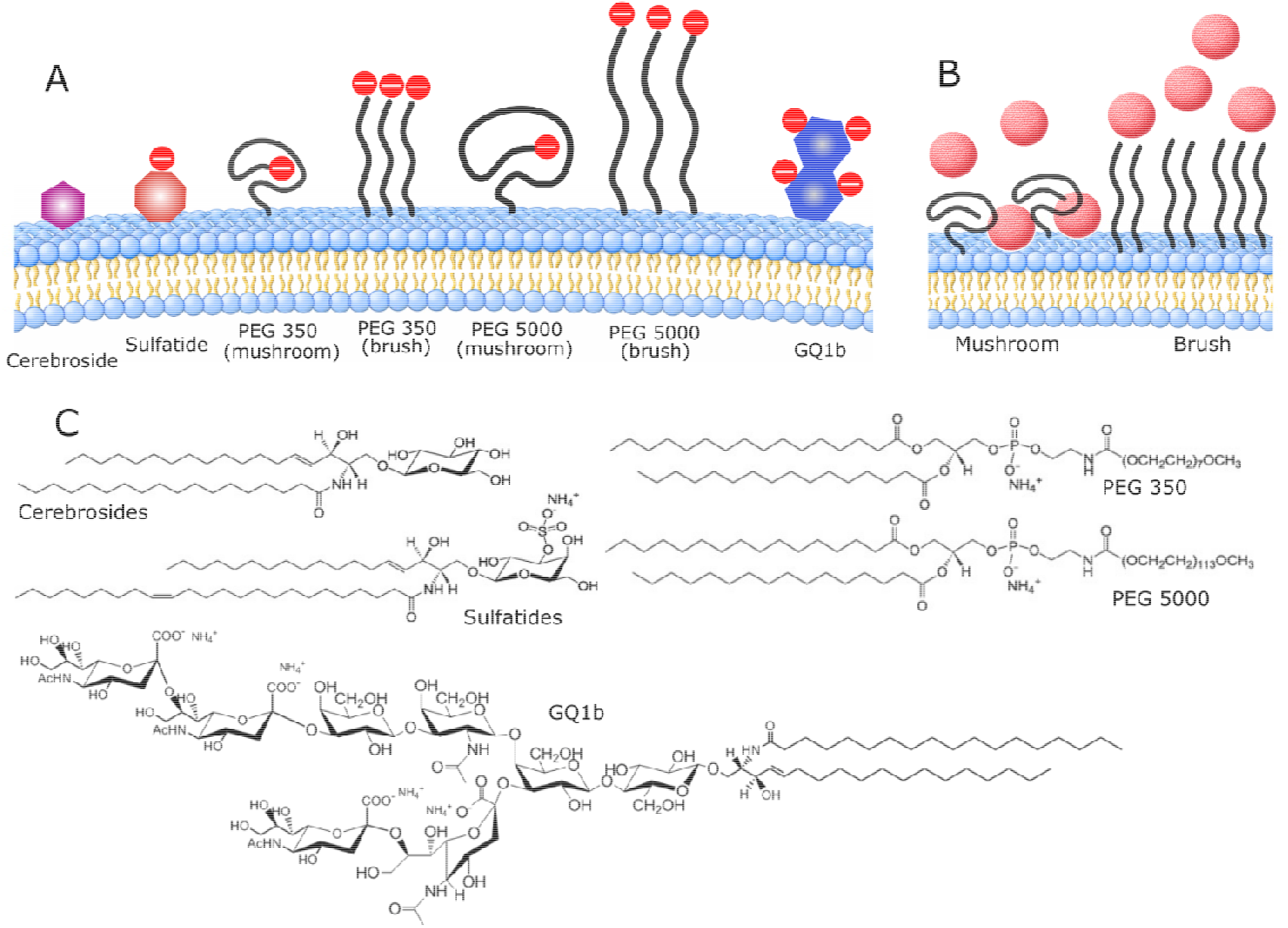
Reconstitution of a glycocalyx mimetic in model membranes. A, the different glycolipids used relative to their sizes and charge. The sketch also shows the folded mushroom and extended brush regimes for the two PEG glycolipids used here. B, expected interactions of liposomes with membranes containing PEG glycolipids in the dispersed mushroom and in the more crowded and extended brush regime. C, the molecular structures of the glycolipids used here.

### Glycolipids do not modify membrane phase state or viscosity

We first investigated whether the reconstitute model glycolipids interfere with membrane mechanics. To probe whether GUVs of made of POPC:Chol and glycolipids exhibit lateral phase separation at the microscale, we performed 3-dimensional reconstructions of the GUVs. It was reported before that cholesterol induces phase separation in mixtures of low and high melting temperature lipids^28^, whereas large polymers anchored to the membrane inhibit it^29^. Because we have a binary mixture of POPC (a low melting temperature lipid) and cholesterol, in combination with glycolipids or not, we do not expect the membranes to phase separate. Indeed, we did not observe phase separation for GUVs devoid of glycolipids, or for any of the glycolipids at any fraction (Figure 2C and Video S1 for a representative glycolipid). Previous observations of the glycolipid GM1 reconstituted at 5 mol% or higher reported a coexistence of liquid-solid domains in POPC GUVs^30^. The absence of phase separation for any of the glycolipids used at much higher fractions is attributed to the presence of cholesterol. We conclude that the glycolipids used in neutral POPC:chol or negative POPC:POPS:chol membranes do not alter the membrane phase state.

**Figure 2.**
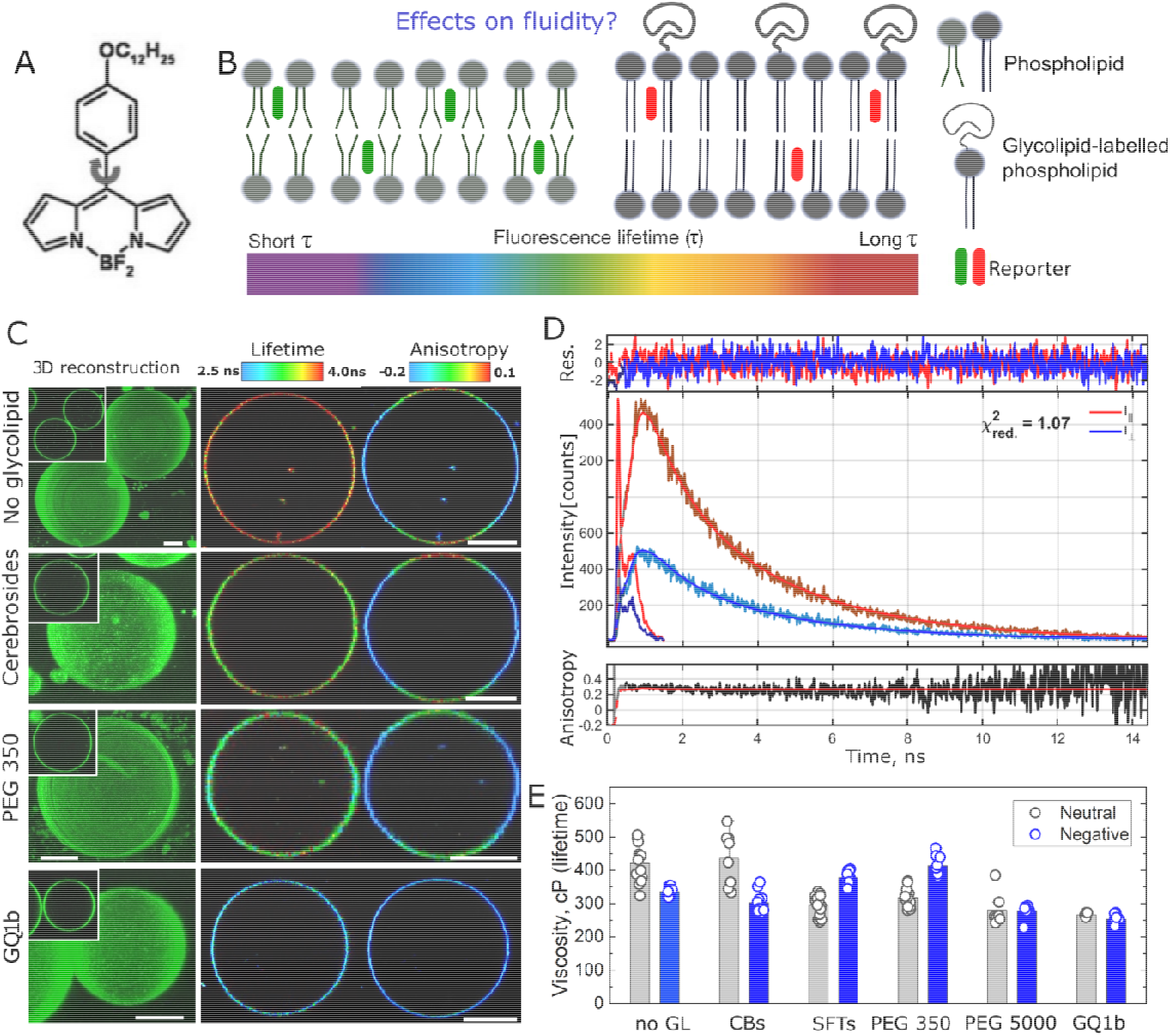
Glycolipids do not significantly alter membrane phase state or viscosity. A, molecular structure of the viscosity-sensitive molecular rotor Bodipy C_12_. B, sketch of viscosity sensing in the membrane. Bodipy C_12_ responds to membrane local viscosity by changes in fluorescence lifetime. Alternatively, changes in the local environment can be also sensed by measuring the probe’s rotation. The unknown effects of glycolipids in the membrane are investigated. C, representative three-dimension (3D) reconstructions of GUVs (left column), FLIM (middle column) and steady state anisotropy (right column) images of negatively charged GUVs containing some of the studied glycolipids. GUVs were made of POPC:POPS:Chol:glycolipid. Scale bars: 10 μm. D, temporal fluorescence intensity decays in the parallel (red) and perpendicular (blue) detector and fits (solid lines). The residuals of the fit for both decays is also shown. Bottom, time-resolved fluorescence anisotropy decay (black) along with its fit (black solid line). E membrane viscosity calculated form FLIM for several individual neutral and negative GUVs containing 20 mol% each of the glycolipids (except for PEG 5000, used at 5 mol%) for neutral (POPC:Chol:GL, grey) and negative (POPC:POPS:Chol:GL, blue) GUVs.

To check whether the glycolipids modify the membrane viscosity, we employed the flexible viscosity-sensitive molecular rotor Bodipy C_12_ as a viscosity reporter (Figure 2A). In fluid media, rotor twisting is relatively unhindered, favouring non-radiative decay, which decreases its fluorescence lifetime^31,32^. In viscous media, twisting is restricted and its fluorescence mainly dissipates radiatively, increasing the fluorescence lifetime. Therefore, by monitoring the fluorescence lifetime, (*τ*), it is possible to assess whether the glycolipids alter the local viscosity (e.g. changes in lifetime map from green to red), using previously calibrated data^31^ (Figure 2B). Alternatively, membrane viscosity was probed by measuring the probe’s molecular rotation. Molecules can rotate freely in fluid media, whereas in more viscous media, rotation in hindered. To assess molecular rotation, we measure Bodipy C_12_ rotational correlation time (*θ*) using time-resolved fluorescence anisotropy^32^. The generated fluorescence decays were collected in two detectors aligned parallel and perpendicular to the excitation, from which *θ* can be used to assess the membrane viscosity (see methods for details).

Figure 2C shows representative FLIM and steady state anisotropy images of GUVs for negative membranes, and Figure S1 show the same data for neutral GUVs. Figure 2D shows the temporal fluorescence intensity decays in the parallel (red) and perpendicular (blue) detectors along with their respective fits and residuals. The figure also shows the time-resolved anisotropy decay and its fit. Figure 2E shows the calculated viscosity obtained from Bodipy C_12_ fluorescence lifetime (molecular twisting) for negative and neutral GUVs reconstituted with various glycolipids. The viscosity from anisotropy measurements (molecular rotation) for these membranes are shown in Figure S2. In general, the local viscosity varied from 300 to 400 cP from lifetime, and from 150 to 250 cP from the anisotropy measurements, in agreement with prior measurements on lipid-only membranes (Lira *et al*., in preparation). Importantly, the values were found to be approximately constant and there was no trend regarding the glycolipids used. From these experiments, we conclude that the glycolipids used here do not significantly modify the membrane viscosity.

### Glycocalyx components alone are sufficient to elicit liposome fusion

We next examined whether glycocalyx components affect the ability of cationic liposomes to bind and fuse with GUV model membranes. Because high amounts of fusion can also be associated with membrane pore formation, we additionally assessed fusion-induced leakage. Notably, the liposomes were fluorescently labelled, hence allowing to resolve liposome docking onto the GUVs.

GUVs were reconstituted with 20 mol% of each of the glycolipids (except PEG 5000, reconstituted at 5 mol%) on their membranes, and incubated with DOTAP:DOPE (50:50, molar fraction) small (∼100 nm size) cationic liposomes. Fusion and leakage were quantified using confocal microscopy from the transfer of the membrane fluorescent probe from the liposomes to the GUVs (fusion), and from the entry of a small aqueous fluorescent probe present in the outer medium (leakage), as sketched in Figure 3A. Leakage was quantified using the degree of filling approach^33^ (see methods for details). In this approach, the degree of filing values vary between 0 and 1; 0 meaning non-permeable vesicles, 1 meaning fully permeable vesicles, and intermediate values representing intermediate degrees of permeability. Fusion also increases the GUV area, inducing visible fluctuations in the GUV membrane^24^. Given their small dimensions^34^, the small liposomes carry more lipids in their outer than their inner leaflets^35^, and this asymmetry is brought to the GUVs via fusion, inducing outward budding^24,36^. By contrast, if hemifusion is the predominant fusion mode, the asymmetric lipid incorporation increases the surface tension, eventually leading to GUV collapse^24^. Therefore, alterations of GUV morphology and structure can additionally be used to probe the fusion mode and efficiency.

**Figure 3.**
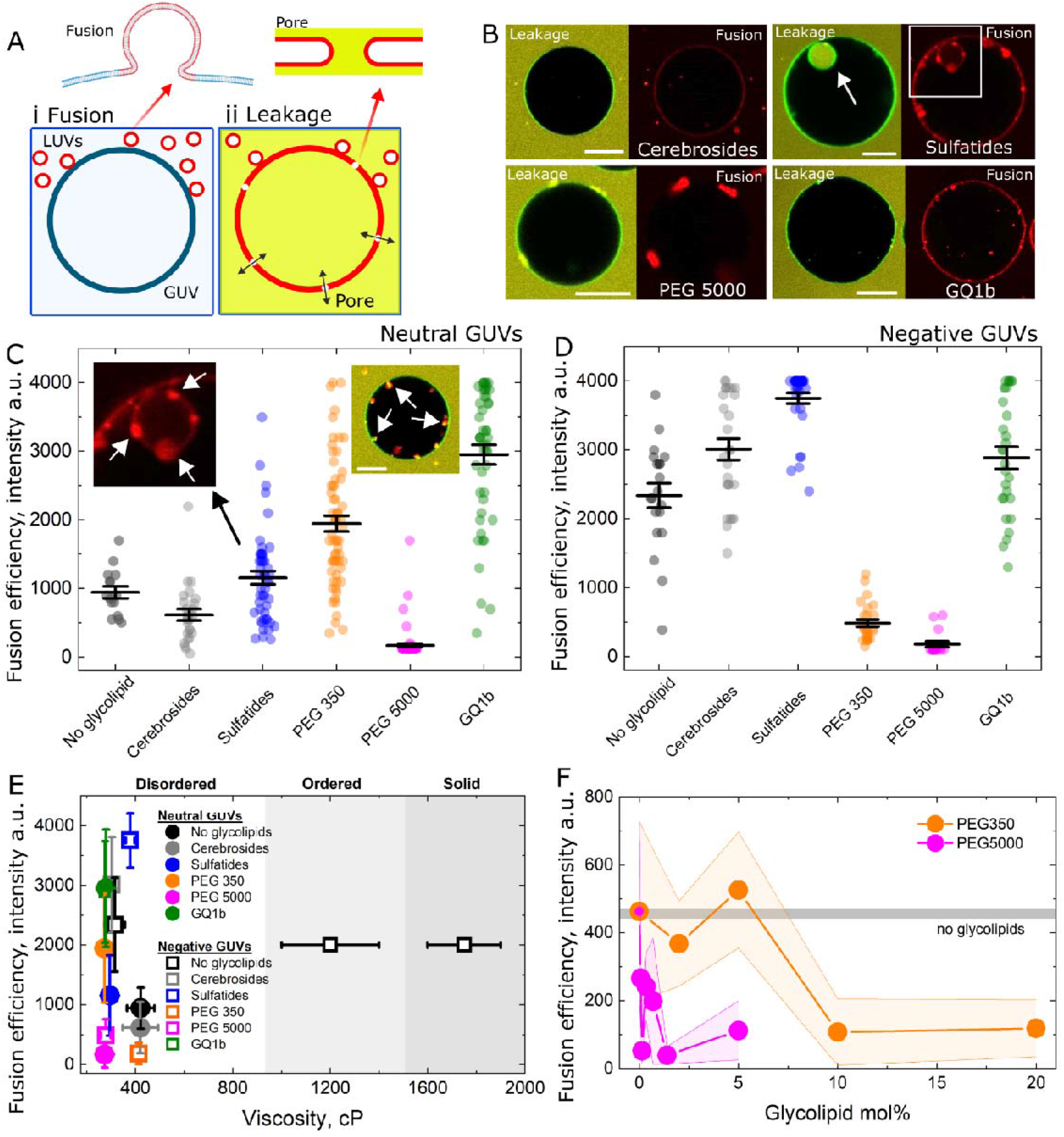
Reconstituted glycolipids interfere with membrane fusion. A, sketch of the fusion assay, wherein membrane-labelled liposomes (red) fuse with GUVs (left, i) and may allow permeation through the GUV (right, ii) due to pore formation (inset). B, representative GUVs reconstituted with some of the glycolipids used in the study and incubated with 10 μM (lipid concentration) liposomes. Leakage and fusion signals are shown. The arrow indicated endocytosis-like budding. Scale bar: 20 μm. C and D, fusion efficiency (lipid mixing) measured for several neutral (in C) and negative (in D) GUVs reconstituted with the various glycolipids used. The GUVs were incubated with 10 μM liposomes (lipid concentration). Means and s.e.m are shown. Inset in C: various liposomes entrapped in endocytosis-like buds for a GUV reconstituted with PEG 5000. E, fusion efficiency as a function of membrane viscosity. The GUVs used here are in the Ld phase. As a reference, the viscosities of GUVs in the ordered and solid phases from a previous report^25^ are shown. F, fusion efficiency as a function of the mol% of PEG 350 (orange) or PEG 5000 (pink). The horizontal grey line represents the efficiency measured on bare (no glycolipid) GUVs. All experiments were performed under identical conditions.

Figure 3B shows representative neutral GUVs reconstituted with different glycolipids, with the neutral GUVs chosen to emulate the outer surface of the plasma membrane. As expected, uncharged cerebrosides did not promote efficient docking or fusion. In contrast, liposomes tended to dock at the surface of GUVs containing PEG5000 but failed to fuse (Figure 3B and Video S2). Liposomes that docked remained mobile on the GUV surface (Video S3). However, liposomes fused efficiently with GUVs reconstituted with the small charged sulfatides or with GUVs with the larger multicharged GQ1b. For these instances of efficient fusion, the GUVs exhibited floppy and fluctuating membranes, a clear indication of fusion-mediated area increase (Video S4). Quantification for all glycolipids is shown in Figure 3C. Again, CBs did not have a significant effect, whereas SFTs enhanced fusion in a subset of vesicles. On average, PEG350 and especially GQ1b significantly promoted liposome fusion with neutral GUVs, while PEG5000 suppressed even the low fusion levels observed in bare neutral membranes.

As a reference, we also examined fusion with negatively charged GUVs containing the same glycolipids, which, as expected, fused much more efficiently than the neutral vesicles (Figure 3D). Again, CBs had little effect, whereas SFTs mildly enhanced fusion, similar to their effect in neutral GUVs. Interestingly, unlike its behavior in neutral membranes, PEG350 strongly inhibited fusion in negatively charged GUVs, with an even stronger suppression observed for PEG5000. This shows that, whereas charged glycolipids promote liposome fusion with neutral membranes, large polymers inhibit fusion even in highly fusogenic conditions. This shows a double effect of glycolipid charge and steric hindrance on liposome fusion. By contrast, fusion with GQ1b-decorated GUVs remained comparable to bare negatively charged GUVs as these were already highly charged. Notably, neutral GUVs containing GQ1b reached similarly high fusion levels to negative bare GUVs. Thus, charged glycocalyx components—particularly multivalent species—can promote fusion of cationic liposomes with otherwise neutral membranes to levels comparable to negatively charged GUVs. Although the representative GUVs in 3B were not permeable, Figure S3 shows that high extents of fusion are associated with high propensity to membrane permeabilization with degree of fillings that scales with fusion; from ∼0.1 for compositions that did not favour fusion (e.g. CBs and PEG 5000), to 0.2-0.6 for conditions that favour fusion (PEG 350 and GQ1b). Such effects did not depend on GUV size Figure S3. These results show that the glycocalyx does not protect the GUVs against permeabilization.

Figure 3E shows the fusion efficiency (measured from the fluorescence intensity) of the different glycolipid-reconstituted GUVs as a function of membrane viscosity (measured from the fluorescence lifetime). All GUVs tested under these conditions remained in the liquid-disordered state. For comparison, the viscosity values previously reported for charged GUVs in the liquid-ordered and solid-ordered states from previously published data are also shown^25^. Previous studies reported that increasing membrane viscosity inhibits membrane fusion^25,37^, but here, all glycolipid-containing GUVs cluster within a narrow viscosity range despite exhibiting a broad spectrum of fusion efficiencies. The results show that cationic liposomes fuse much less efficiently with neutral than with negative GUVs. A notable exception is neutral GUVs reconstituted with GQ1b, which promoted fusion to levels comparable to negatively charged GUVs, as long as these were not decorated with large polymers that inhibit fusion. Because these large differences in fusion (from intensity values 200-4000 a.u.) occurred with minor changes in membrane viscosity (∼ 300-400 cP), we conclude that glycolipids alone are sufficient to regulate liposome fusion with cell-like GUVs independently of membrane mechanics.

To probe how liposome fusion is regulated by the polymer conformation, we added cationic liposomes to negatively charged GUVs that were coated with increasing molar fractions of either PEG350 or PEG5000. At low surface coverage, flexible PEG chains adopt isolated coil-like conformations, the so-called mushroom regime, whereas at higher coverage, neighbouring chains begin to overlap and extend away from the membrane, forming a denser brush regime^27^. In addition to surface density, the mushroom-to-brush transition strongly depends on polymer molecular weight, and is expected to occur at ∼11 mol% for PEG350 and ∼0.5 mol% for PEG5000^27^. As shown in Figure 3F, fusion was progressively inhibited with increasing grafting density for both polymers, which happened at different concentration ranges. Relative to the control without polymer (horizontal grey line), PEG350 did not measurably reduce fusion up to 5 mol%, whereas fractions of 10 mol% and above completely abolished it. PEG5000 also fully suppresses fusion, but at much lower fractions as only 0.05 mol% already significantly reduced fusion while 2 mol% was sufficient for complete inhibition. Because fusion was abolished once the polymers enter the brush regime, these results indicate that polymer conformation plays an important role in regulating liposome fusion with polymer-decorated GUVs, either by directly interfering with fusion itself, or by sterically preventing liposome binding to the membrane.

### An artificial glycocalyx regulates liposome fusion *vs*. endocytosis

GUVs dispersed in glucose solutions tend to bend inward in an endocytosis-like process, forming large buds of a few micrometer diameter, irrespective of the presence or identity of glycolipids or interaction with liposomes (Videos S5-S6). This is in agreement with prior observations for membranes devoid of glycolipids^38,39^, and a consequence of the spontaneous curvature induced by different sugar solutions across the GUV membrane^40^. Because the magnitude of curvature is small, the formed buds are of micrometre dimensions. This endocytosis-like process entraps solutes from the medium (e.g. fluorescent dyes; see Figure 3B for the SFTs GUV). Fluorescence recovery after photobleaching (FRAP) shows that the endocytosed bud remains connected to the main GUV membranes, with a bud neck that, in some GUVs, can be wide enough to allow small ‘endocytosed’ cargo to exchange with the outside medium (Figure S4 and Video S7), whereas in some other GUVs, the bud neck is too tight to allow content exchange (Video S8). For conditions of very efficient fusion, liposome binding was immediately converted to fusion. In contrast, in less efficient fusion conditions, the liposomes remained largely bound to the membrane and became passively internalized in these “endocytic” structures; e.g. sulfatides in conditions of fusion, inset in Fig. 5C, and PEG 5000, inset in Fig. 5C, when fusion is inhibited. Note that solutes from the medium are co-internalized in the process (i.e. yellow signal from SRB in the buds). FRAP on these internalized liposomes show that the liposomes remain trapped in the buds (Figure S4 and Video S9). Because efficient liposome fusion induces outward rather than inward budding, endocytosis produces GUVs that simultaneously exhibit inward and outward buds (Video S10).

Upon GUV incubation with liposomes, the generated buds did not only entrap liposomes bound to the GUV surface, but under conditions of efficient fusion, the buds themselves underwent extensive remodelling, generating tubes that protrude away from the bud (towards the GUV lumen). These tubules formed from vesicles that either remained intact (Figure 4C below) or became permeable (Figure S5), indicating that pore formation is not required for their generation. The resulting tubes were highly dynamic, exhibiting thermal fluctuations and bending, entrapping small solutes from the medium (Figure 4D below) and they were generated in GUVs decorated with glycolipids that enable liposome fusion, such as sulfatides or GQ1b (Figure S6). FRAP measurements showed rapid fluorescence recovery (Video S11), demonstrating that the tubes remain continuous with the bud membrane, which itself remains connected to the parent GUV. We hypothesize that the tube formation originates from curvature generation driven by an imbalanced distribution of glycolipids following fusion. Fusion with liposomes dilutes the glycolipids in both leaflets of the GUV membrane; however, because the fusing liposomes contain an excess of outer leaflet lipids, dilution of glycolipids in the outer leaflet of the GUVs is disproportionately greater, generating an area mismatch that bends the membrane toward the side with the excess area.

**Figure 4.**
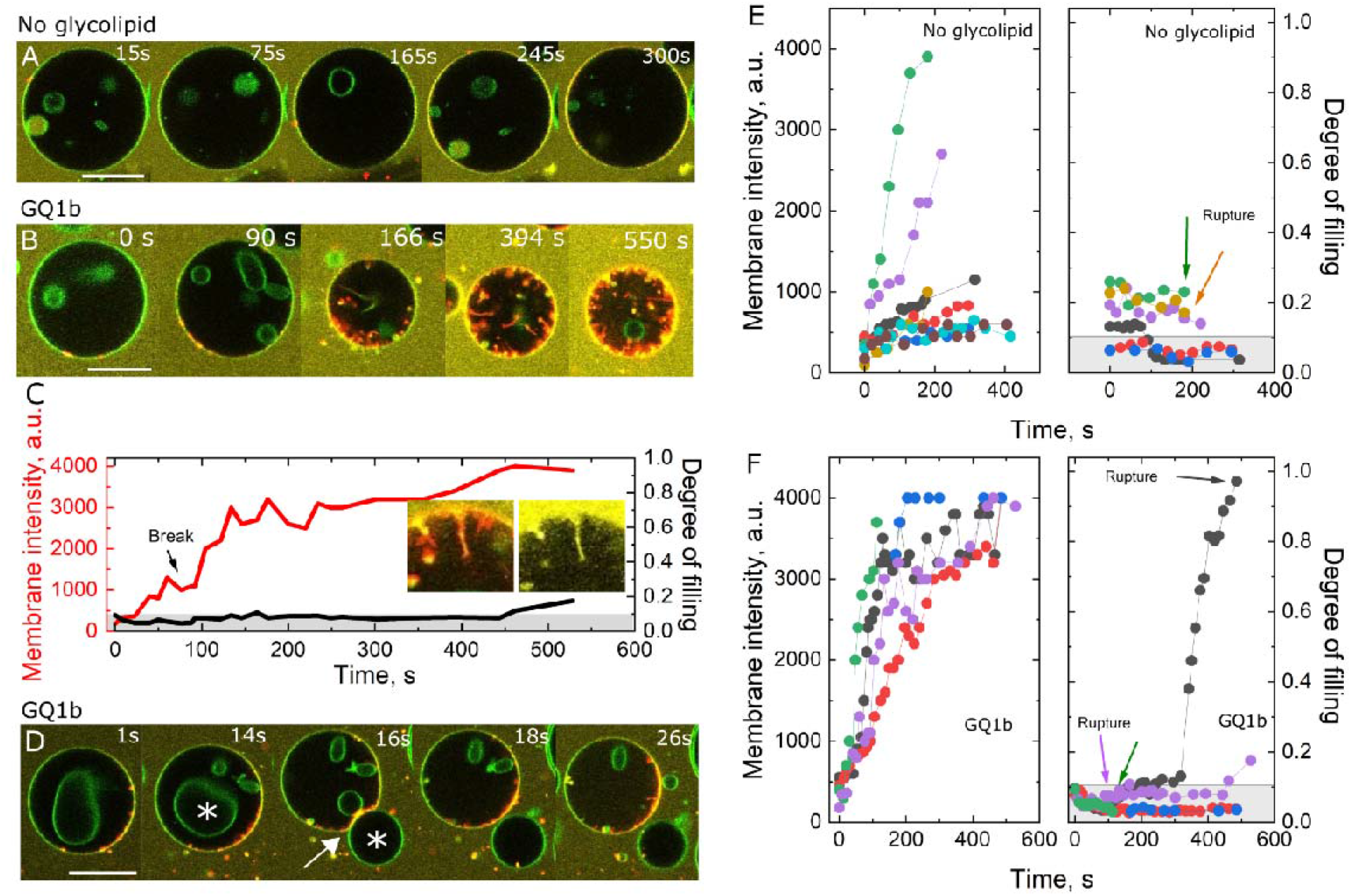
Cationic liposomes undergo slow hemifusion with GUVs decorated with an artificial glycocalyx. A, representative snapshots of cationic liposomes (red) interacting with a neutral GUV (green) in the presence of SRB (yellow). B, rupture and inward tube formation of a GQ1b-decorated GUV upon liposome fusion. C, dynamics of fusion as assessed from the increase in GUV membrane intensity and leakage, as assessed form the degree of filling. Times zero correspond to the beginning of observation. D, rupture of a GUV, enabling the scape of an inner vesicle (asterisk) through a pore (arrow) at time 16 s. The Scale bar: 20 μm. The cationic liposomes are made of DOTAP:DOPE (50:50, molar ratio) and labelled in red. The GUVs are made of POPC:Chol (7:3, molar ratio) or POPC:Chol:GQ1b (5:3:2, molar ratio) and labelled with 0.2 mol% Bodipy C_12_ (green). SRB was present in the medium at 10 μM. E and F, fusion (left) and leakage assessed from the degree of filling method (right) for neutral GUVs devoid of glycolipids (in E) or containing GQ1b (in F). Rupture events are indicated by an arrow. The grey bar represents non-permeable GUVs.

In addition to our recent findings that molecules capable of flip flop can remodel growing membranes^41^, we here thus show that glycolipid-decorated membranes also undergo extensive remodeling upon growth. Irrespective of the generation mechanisms, the results show a new type of membrane remodelling associated with glycocalyx-dependent liposome fusion and curvature generation.

### A charged artificial glycocalyx promotes fusion via a slow hemifusion intermediate

We locally injected liposomes in the vicinity of neutral GUVs containing GQ1b as a model glycocalyx to follow the fusion and the associated remodelling processes in real time. The results are shown in Figure 4E for the dynamics of the fusion and permeability for several GUVs, and for comparison for a representative bare GUV in Figure 4A. We observed that liposomes hardly fused with neutral and bare GUVs. Note liposome docking on the GUV surface. The degree of filling data in Figure 4E also indicates some rupture events, identified as sudden GUV jumps due to pore formation and expulsion of inner content^42^. Although some extent of leakage was observed, this remained modest.

When we performed these experiments with GUVs decorated with GQ1b, however, we observed that the liposomes docked very efficiently on these GUVs, which is not surprising given the high charge of GQ1b, which resulted in greater fusion compared to bare GUV without glycolipids. However, liposome fusion was slower than fusion with negatively charged GUVs, where docking was almost instantaneous, followed by fusion within sub-second timescales^43^. Because docking increases surface tension^44,45^, it is not unexpected that these GUVs rupture, but due to the high edge tension in POPC:Chol GUVs^46^, pores formed during fusion eventually closed, with only a small extent of leakage observed. This is clearly outlined in Figure 5B (see also Video S12), where rupture noticeably decreased GUV size due to volume loss albeit with minimal leakage. As shown in the dynamics in Figure 5C, docking did proceed to fusion. From these sequences, it is possible to see the generation of a large number of inward tubes produced in the GUV as fusion progressed. Note that these tubes did also entrap the aqueous content, showing that their interior was continuous with the outer medium. Figure S5 shows inward tubulation and collapse for a GUV that became permeable. For a GUV that contained a vesicle in its interior (which is common in GUVs prepared by electroformation), the docking-mediated increase in tension opened a pore large enough to allow the expulsion of an inner vesicle (Figure 5D; see also Video S13). Again, however, because of the limited pore lifetime, the leakage was limited. Compared to bare GUVs, the liposomes fused more efficiently when the GUVs were decorated with GQ1b.

**Figure 5.**
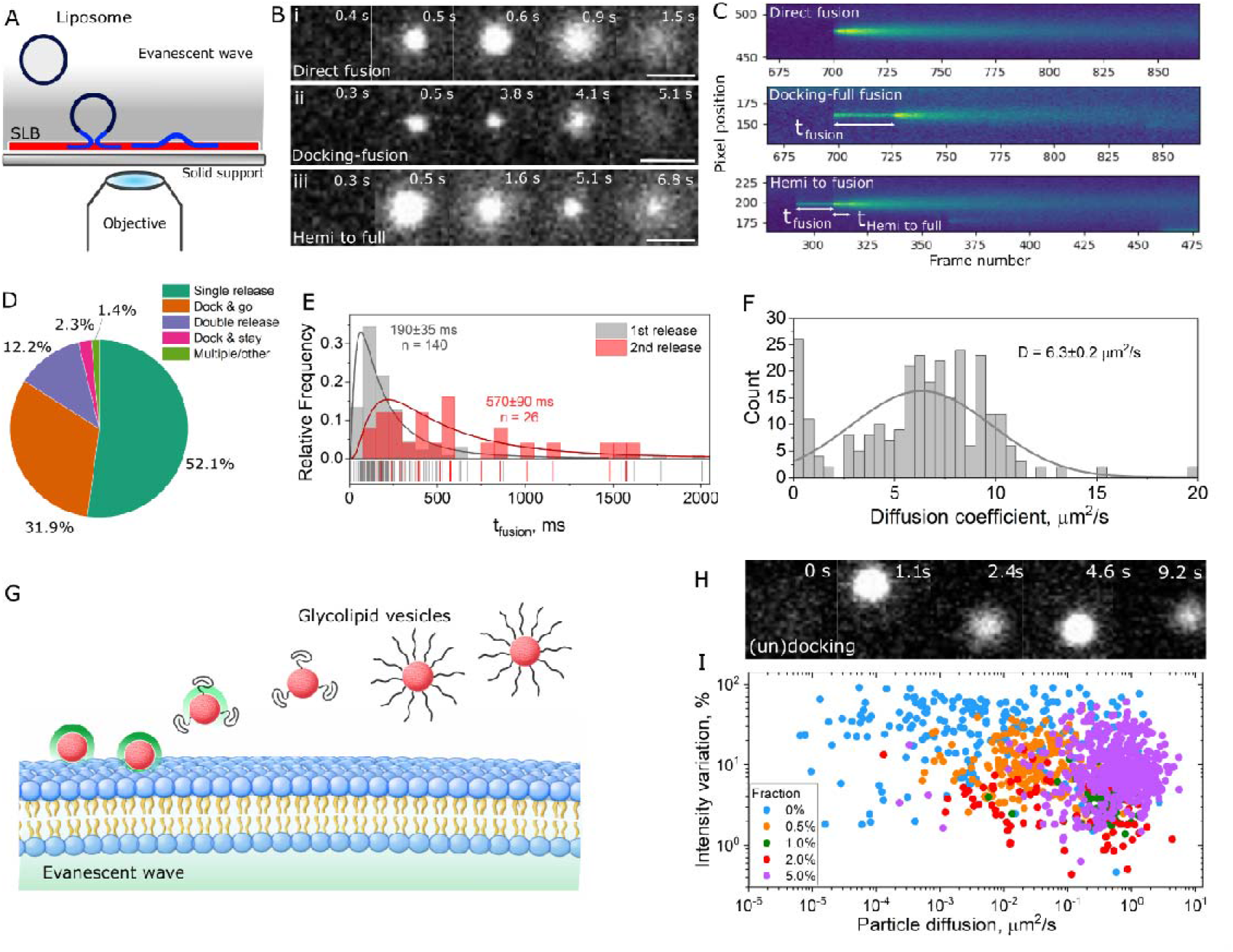
Large polymers mainly interfere with the docking stage. Sketch of the single vesicle fusion assay. Incoming fluorescent liposomes above the evanescent field are not visible (indicated as black), whereas approaching to the supported lipid bilayer (SLB) results in the appearance of a bright diffraction-limited spot. Fusion brings the labelled liposome closer to the surface, increasing fluorescence intensity (indicated as blue). B, representative snapshots of single liposomes interacting with the SLB. i, ii and iii show direct fusion, docking to fusion transition and hemifusion to full-fusion transition for bare liposomes devoid of glycolipids. Scale bars: 2 μm. C, kymographs of the events shown in B. The fusion transitions and their corresponding times are also indicated. D, pie chart representing the fractions of each observation. E, histogram distribution of single (1st release) and double (2^nd^ release) fluorescence release events, representing direct fusion, and hemifusion followed by full fusion, respectively. The mean and s.d. are shown. F, measured diffusion coefficients for incoming lipids resulting from liposome fusion to the SLB. The mean and s.d. are shown. G, sketch of the interactions of polymer-grafted liposomes. Close proximity to the bilayer results in an increase in fluorescence signal (indicated in green) due to TIRF excitation. Note the mushroom and brush conformations. H, representative docking of a liposome containing 2 mol% PEG 5000 that repeatedly docks (sharp spot) and undocks (dim spot) the SLB. I, 2-dimensional plot of fluorescence intensity variation within the TIRF excitation as a function of lateral diffusion for liposomes with varying fractions of PEG 5000. Each dot represents a measurement.

Because the fusion of large numbers of liposomes increases the area of neutral GUVs decorated an artificial glycocalyx, as evidenced by appearance of membrane fluctuations and outward budding, it is clear that the liposomes undergo full fusion with the GUVs (since mere docking or hemifusion do not lead to increase in membrane area)^24,26^. However, since the liposomes remained docked for minutes - in contrast to fast fusion with negatively charged GUVs - the tension increased, eventually rupturing the GUV membrane. These docked liposomes proceeded through a slow hemifusion intermediate, undergoing lipid mixing with the GUVs (i.e. increase in red fluorescence, as shown in Figs. 4B,F) without an immediate area increase. Thus, although the glycocalyx-mimetic system promotes fusion between cationic liposomes and otherwise neutral synthetic membranes, the process transits through a long-lived hemifusion state.

### Steric hindrance inhibits liposome docking

The liposome–GUV experiments demonstrated that charged glycolipids can convert otherwise neutral, docking-prone GUVs into fusion-competent membranes through a slow hemifusion intermediate. However, while charged glycolipids promote fusion, large polymers can suppress it even under highly fusogenic conditions because of steric hindrance. Although the GUV assay robustly captures overall fusion outcomes, it did not distinguish whether this steric inhibition arises from impaired liposome binding or from fusion following binding.

To resolve these mechanisms, we monitored liposome interactions with solid-supported lipid bilayers (SLBs) at the single-particle level using high-speed total internal reflection fluorescence (TIRF) microscopy. SLBs composed of the cationic membrane were incubated with negative bare liposomes lacking glycolipids or containing increasing fractions of PEG 5000. In this assay, liposome docking appears as a bright diffraction-limited spot within the evanescent field, whereas its sudden disappearance (labelled as “go”) indicates unbinding^47^ (Figure 5 A,G). Fusion is identified by a rapid fluorescence burst as labelled lipids enter the evanescent field upon the membrane merger, followed by an exponential decay as lipids diffuse away from the fusion site^48^. Persistent post-burst fluorescence reported hemifusion, whereas a full decay to background indicated full fusion, allowing direct resolution of fusion intermediates and kinetics^49^. This assay thus directly separates steric effects on liposome docking from inhibition of downstream fusion.

The formed SLB is fluid and has a measured lipid diffusion coefficient of 0.8 μm^2^/s (Figure S7), which is comparable to other SLB studies^50,51^ and one order of magnitude slower than that observed for free-standing membranes^52^, due to friction with the underlying support^43^. Figure 5B shows representative fusion results. In i, a liposome docked and immediately fused with the SLB within one frame. Because there is no remaining liposomal signal, this indicates direct full fusion. In ii, the liposome docked to the SLB for a few seconds before full fusion occurred. In iii, the liposome immediately fused with the SLB, but it left a permanent bright spot behind, before a second fluorescence burst occurred that decayed to ∼ background. The process can be ascribed to hemifusion followed by fusion of the inner leaflets. These three events are shown in Movies S14-16 and were observed with liposomes devoid of glycolipids.

To quantitate these events, we transformed the fusion traces into kymographs (Figure 5 C), from which each fusion intermediate, fusion kinetics and diffusion coefficient of the laterally diffusion lipids can be determined (see Figure S8 and methods for details). Upon binding to the SLB, bare liposomes did not diffuse laterally in the membrane, which is evidence of strong interactions. As shown in Figure 5D, from ∼500 single liposomes analysed, most (52%) bare liposomes underwent direct fusion, resulting a single release event, and only a few (12%) underwent a resolvable hemifusion step (first release) before full fusion (second release). A fraction of the docked liposomes (32%) disappeared (dock & go), which can either indicate weak binding or occasional and random approach to the SLB within the TIRF excitation field and diffusing away. Approximately 2% of the liposomes stably bound to the SLB without undergoing fusion within the experimentally observed time (Dock & stay). For a small fraction ∼ 1%, we detected multiple releases, which are either a result of fusion of multilamellar liposomes, or fusion of multiple liposomes in close proximity below the optical resolution. These were excluded from further analysis. We also quantified the kinetics of transition from the fusion intermediates. As shown in Figure 5E, most liposomes fuse in a time < 200 ms. For a small and resolvable fraction, the transition from the first to the second release event (the time from hemifusion to full fusion) was resolved at an average time of ∼ 0.6 s.

Notably, these events are among the fastest single events reported, especially compared to protein-mediated vesicle fusion, which exhibit fusion times in the order of tens of seconds^49,53,54^. For proteoliposomes, fusion seems to be more efficient for smaller vesicles due to the high tension in highly curved membranes^55,56^. For the charged liposomes used here, neither direct fusion nor the transition from hemi to full fusion depended on particle size (Figure S9). From the lateral diffusion of lipids from the fusing liposomes to the SLBs, we calculated a diffusion coefficient as ∼6 μm^2^/s using eq. 2, which is significantly higher than that measured on prepared SLBs.

We next checked whether liposome binding and fusion are affected by a glycocalyx-like polymer mesh on the liposomes. From the liposome-GUV experiments, we expected that an increasing fraction of PEG 5000 would change the liposome behaviour from strongly interacting to less interacting and non-fusing liposomes, as sketched in Figure 5G for polymers in the mushroom and brush regimes. Fluorescence correlation spectroscopy (FCS) measurements of the three-dimensional diffusion coefficient (D) of liposomes in solution grafted with increasing PEG 5000 surface coverage, showed a mild decreases from 2.5 to 1.7 μm^2^/s as PEG 5000 increased from 0-0.75 mol%, sharply decreasing to a constant value of D ≈ 1.0 μm^2^/s beyond 1 mol% (Figure S10). This can be ascribed to the experimentally observed mushroom-to-brush transition regime observed for PEG 5000 at 0.5 mol%^27^.

In contrast to bare liposomes, which bind firmly to the SLBs and do not move laterally, liposomes containing 2 mol% PEG 5000 (i.e. in the brush regime) were found to bind weakly to the SLB and they did not fuse (Figure 5H and Video S17). Instead, they underwent lateral motion and occasional detachment. For a liposome in the brush regime diffusing and D = 1 μm^2^/s, the mean square displacement in the axial direction is <z^2^> = 2Dt. Considering a TIRF penetration depth of 150 nm^47^, a freely diffusing liposome in this regime that does not interact with the membrane stays in the field of view for approximately one frame (33 ms/frame). This sets the time for random appearance in the TIRF field of view. From single liposome fusion experiments with PEG 5000 liposomes Random appearance is the case of ≈ 20% (23 out of 105) vesicles containing 2 mol% PEG 5000 and ≈ 30% (63 out of 198) of vesicles containing 20 mol% PEG. This means that the remaining 70-80% of liposomes do interact with the underlying bilayer (residence time > 1 frame), albeit weakly, but this interaction does not progress to fusion. Furthermore, only 2% (2 out of 105) of the docked liposomes containing 2 mol% PEG 5000 underwent hemifusion (but not full fusion) and these hemifusion times were in the order of 300 ms, slower than hemifusion-to-full fusion events for bare liposomes. We did not observe any (hemi or full) fusion event for liposomes containing higher PEG 5000 fractions. We conclude that PEG 5000 glycolipids impairs liposome binding and fusion.

To quantitatively assess how the PEG 5000 mol fraction affects liposome–SLB engagement, we measured the percentage variation in particle fluorescence intensity during the observation window together with the corresponding lateral diffusion during interaction. The fluorescence-intensity variation serves as a discriminatory parameter between liposome fusion and simple membrane association: fusion leads to two-dimensional spreading of lipids into the SLB, resulting in a pronounced decrease in particle fluorescence intensity, whereas membrane-bound liposomes that diffuse laterally without fusing retain a largely constant intensity. An informative way to interpret these data is a 2D phase space of liposome–SLB interaction states where data are plotted as intensity variation versus the diffusion coefficient (Figure 5I). High D values indicate fast lateral motion, weak confinement, and transient or absent binding. By contrast, liposomes that exhibit engagement and subsequent dye release exhibit a high intensity variation, as they essentially disappear. This is clearly observed with the 0% PEG data in Figure 5I. Increasing PEG5000 densities progressively shift the liposome population from an engaged state toward a freely diffusing, non-interacting state. The results corroborate that PEG 5000 suppresses the formation of a stable fusion-competent docking state, thereby preventing fusion. Rare liposomes that do bind at low polymer fractions either fail to fuse or do so with slower kinetics.

## Conclusions

Under physiological conditions, the outer leaflet of the plasma membrane contains only low fractions of anionic phospholipids, and electrostatic interactions with the cell surface arise predominantly from glycocalyx components rather than from membrane lipids themselves^2,5,7^. Previous experiments with GUV model membranes showed that fusion of cationic liposomes requires anionic lipids, whereas neutral GUVs mainly support docking. The viscosity of the outer leaflet is also high compared to the inner leaflet^11^, and an increase in viscosity further inhibits fusion^12,25^. In these synthetic systems, efficient fusion increases the membrane area and alters spontaneous curvature^36^, while docking and stalled hemifusion instead raise membrane tension which can ultimately drive vesicle rupture^24^. Strikingly, the same liposomes fuse efficiently with living cells^23^, despite the absence of significant charged lipids at the cell surface. This discrepancy reveals that minimal synthetic membranes lack key interfacial components present in natural membranes that function as charge receptors. Here, we identified the glycocalyx as this missing component that bridges the gap between synthetic and biological membranes.

Introducing highly charged glycocalyx components provides an electrostatic force for liposome binding and fusion, although the outcome is nuanced because glycolipids simultaneously impose steric constraints. For moderately sized, highly charged polymers, we found that liposomes fully fuse with synthetic membranes, increasing membrane area and remodeling curvature. However, fusion was observed to proceed more slowly than with bare negatively charged membranes and to transit through a long-lived hemifusion intermediate that tends to increase membrane tension. This slower pathway likely reflects a reorganization of the polymer mesh to permit close membrane binding, preparing the membranes for fusion. For larger polymers, steric hindrance dominates and fusion is blocked altogether. Single-vesicle measurements show that this inhibition occurs primarily at the docking step, preventing liposomes from establishing a stable fusion-competent contact. And even upon docking, fusion proved to be inefficient. These effects depend strongly on the grafting density and polymer conformation, being most pronounced in the extended brush regime (sketched in Fig. 1B), and they occur independently of membrane viscosity. Under selected conditions, the same glycocalyx components also promoted endocytic budding, enabling internalization of molecules and particles from the surrounding medium. Particle uptake occurred under both fusogenic and non-fusogenic conditions, and the captured cargo remained stably trapped within the endocytic bud, despite its continuous membrane connection to the parental vesicle.

These findings establish the glycocalyx as a fundamental regulator of membrane interactions, particularly for fusogenic particles such as enveloped viruses, extracellular vesicles, pathogen-associated nanoparticles and drug- and gene-delivery systems. Because high expression levels of glycocalyx components are associated with increased metastatic outcome in cancer cells^57^, they represent an enormous potential for cationic drug-delivery systems. More broadly, the findings reveal that the glycocalyx is not merely a passive surface decoration, but a fundamental interfacial layer that governs how membranes sense, capture, repel, and remodel upon contact with external objects. As such, the rational design of synthetic cells that robustly communicate and interact with their environment will inevitably require the engineering of glycocalyx-like systems.

## Materials and Methods

All materials and chemicals were used as obtained. Glucose, sucrose and the fluorescent dye sulforhodamine B (SRB), were purchased from Sigma-Aldrich (St. Louis, MO, USA). The glycolipids brain cerebrosides, brain sulfatides, 1,2-dioleoyl-sn-glycero-3-phosphoethanolamine-N-[methoxy(polyethylene glycol)-350] (ammonium salt) (PEG-350), 1,2-dioleoyl-sn-glycero-3-phosphoethanolamine-N-[methoxy(polyethylene glycol)-5000] (ammonium salt) (PEG-5000), Ganglioside GQ1b (Porcine Brain), the phospholipids1-palmitoyl-2-oleoyl-sn-glycero-3-phosphocholine (POPC), 1-palmitoyl-2-oleoyl-sn-glycero-3-phospho-L-serine (sodium salt) (POPS), 1,2-dioleoyl-sn-glycero-3-phosphocholine (DOPC), 1,2-dioleoyl-sn-glycero-3-phospho-(1’-rac-glycerol) (sodium salt) (DOPG), the lipid cholesterol and the fluorescent probe 1,2-dioleoyl-sn-glycero-3-phosphoethanolamine-N-(lissamine rhodamine B sulfonyl) (ammonium salt) DOPE-Rh were purchased from Avanti Polar Lipids (Alabaster, AL). The probe DOPE-Atto 655 was purchased from Atto-Tec (Siegen, Germany). The molecular rotor Bodipy C_12_ was kindly donated by Klaus Suhling (King’s College London).

Small unilamellar vesicles (SUVs) were prepared by the hydration method followed by bath sonication. Shortly, a 2 mM lipid solution containing the respective lipids were dissolved in chloroform under N_2_ atmosphere and stored at -20C before use. For SUV preparation, the lipid solution was placed in a glass vial and chloroform was evaporated under a stream of Argon. If not stated otherwise, the SUVs were made of DOTAP:DOPE (50:48 molar ratio) and labelled with 2 mol% DOPE-Rh. Chloroform residues were further removed for ∼ 2h under vacuum, forming a dry lipid film. The lipid film was hydrated with a 200 mM sucrose solution, vigorously vortexed until full detachment and extruded 11 times through a 100 nm polycarbonate filter with the help of a manual extruder (Avanti Polar Lipids, Alabaster, AL). The typical stock lipid concentration was 2 mM, and when required, the solution was diluted in the same sucrose solution. The formed SUVs were stored at 4°C and used withing 2-3 weeks.

The GUVs were prepared by the electroformation method^58^ with minor modifications^42^. Briefly, 10-20 ml volume of a lipid solution (2 mM) in chloroform was spread on the surface of two indium titanium oxide slides and evaporated under a stream of nitrogen. The slides were sandwiched with the help of a Teflon spacer, creating a volume of ∼ 1.8 ml. The lipid films were hydrated with a 200 mM sucrose solution and connected to a function generator to apply a 5 mV nominal Vpp voltage and a sinusoidal AC current of 10 Hz for 1-2h. Afterwards, the GUVs were harvested for use. Neutral GUVs were made of POPC:Chol:glycolipid (50:20:30:20, mole fraction), and negative GUVs made of POPC:POPS:Chol:glycolipid (30:20:30:20, mole fraction) and labelled with 0.2 mol% Bodipy C_12_.

The GUVs (50 ml) were incubated with the 10 mM liposomes (lipid concentration) in an Eppendorf tube for 10 minutes for a final 100 ml solution (the remaining volume made of isosmolar sucrose). After incubation, the samples were diluted in 300 ml isosmolar glucose for observation. Both incubation and imaging were performed at room temperature. For 3-dimensional imaging, the 50 ml of GUVs were diluted in 350 ml isosmolar glucose and let it rest for ∼ 15 minutes before imaging.

Supported lipid bilayers (SLBs) were formed from liposome rupture on a glass slide as in^43^. A small volume of a concentrated solution of CaCl_2_ was added to bath sonicated cationic liposomes made of DOTAP:DOPE (50:50, molar ratio) and labelled with 0.2 mol% DOPE-Rh for a final concentration of 2 mM. This solution was transferred to an observation chamber coated with a glass slide and incubated at R.T. for 30 minutes. Non-ruptured liposomes were removed from 3 through rinses with an isosmolar sucrose a solution, after which the SLBs are ready. The quality and fluidity of the SLBs were checked from homogeneous fluorescence and FRAP.

Confocal imaging was performed on a Nikon Eclipse Ti confocal microscope using NIS-Elements AR Nikon-software. The samples were imaged through a 60x water-immersion objective (NA: 1.27). Images were acquired with a Galvano High Resonant Scanner at 512×512 pixels in a sequential mode to minimize bleed-through. Bodipy C_12_ was excited with a 488 nm laser line and emission detected with a 500-550 nm band pass filter. DOPE-Rh was excited with a 561 nm laser line and emission detected with a 570-620 band pass filter. For 3-dimensional imaging, z-stacks were performed at a 0.7 mm step size. All the images were analyzed using Fiji (NIH, USA). GUV fluorescence intensity measurements were performed by a computing DOPE-Rh intensity from a line cross section along the GUV equator.

GUV permeability was quantified using the degree-of-filling (DOF) method, which measures the equilibration of a fluorescent solute between the vesicle lumen and the external solution^59^. In brief, GUVs and LUVs were incubated for 10 min in the presence of 10 mM sulforhodamine B, after which the samples were transferred to an observation chamber for imaging. Changes in intravesicular fluorescence intensity report membrane permeability in GUVs as the degree of filling, which can be calculated as

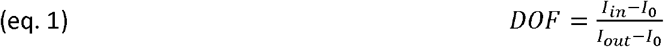

where *I*_in_ and *I*_out_ are the fluorescence intensities measured inside and outside the GUV, respectively, and *I*_0_is the background fluorescence. DOF values range from 0 (impermeable) to 1 (fully permeable). To account for background noise, we pragmatically defined a permeability threshold of 0.1, such that GUVs with DOF values < 0.1 were considered non-permeable.

Fluorescence recovery after photobleaching was performed on the same microscope. For FRAP on SLBs, photobleaching was performed using the argon 561⍰nm laser at maximum intensity for 1 s to bleach DOPE-Atto655 through a circular ROI of nominal radius r_n_⍰= 5⍰μm, and emission detected with a 570-620 band pass filter. The laser was then switched back to attenuated intensity and the recovery images were recorded for 80 s. The fluorescence intensity recovery was used to calculate the lateral diffusion coefficient D. From the recovery curve, D can be calculated from the half-time of fluorescence recovery t_1/2_^60^ (i.e., the time to reach F_1/2_ ⍰=⍰(F_o_⍰+⍰F_∞_)/2, where F_o_ and F_∞_ are the fluorescence intensity in the first post-bleach image and after full recovery), according to

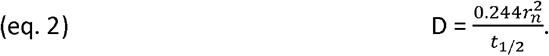

FLIM (Fluorescence lifetime imaging microscopy) and time-resolved fluorescence anisotropy were performed on a Microtime200 microscope (PicoQuant, Germany) using the Symphotime64 software as described previously^61^. Bodipy C_12_ was excited with a 485 nm pulsing laser with a repetition frequency of 40 MHz and 10 µW power as measured at the imaging plane. The emission light was passed through a 50 µm pinhole, split by polarization, and filtered by 525/50 band pass filters (Chroma, USA) before being focused on single-photon avalanche-diode detector (PD5CTC and PD1CTC, Micro Photon Devices, Italy). GUVs were first localized using widefield imaging and then imaged in confocal mode using an objective piezo monodirection scanner, typical 200×200 px size (0.4 μm/pixel) and 0.6 ms pixel dwell times. The data was analyzed in the PAM software package using MATLAB^62^. The bright rim of individual GUVs was selected using a combination of intensity thresholding and manual selection. The intensity decays of the parallel and perpendicular detection channels were globally fitted using a model combining a bi-exponential fluorescence intensity decay with a single-exponential anisotropy decay using the approach of Schaffer et al.^63^.

We used two methods to calculate membrane viscosity (*η*_m_), the first based on fluorescence lifetime, and the second based time-resolved anisotropy. The fluorescence decay of Bodipy C_12_ was fitted using a bi-exponential decay model

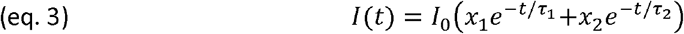

where *I*_0_ is the total fluorescence intensity at t= 0, *τ*_1_ and *τ*_2_ are the fluorescence lifetimes of the two emissive populations, and *x*_1_ and *x*_2_ are their fractional amplitudes, with *x*_1_ + *x*_2_ = 1. The average fluorescence lifetime *τ* was calculated from the fitted parameters and converted into membrane viscosity using a Förster–Hoffmann calibration, in which the logarithm of the lifetime scales linearly with the logarithm of viscosity

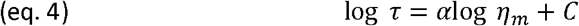

where *α* is the calibration slope and *C* is a constant determined experimentally^64,65^. This relation enabled direct extraction of membrane viscosity from the fluorescence lifetime measured in each pixel of the FLIM images.

Alternatively, *η*_m_ was inferred from time-resolved fluorescence anisotropy. The anisotropy decay was calculated as

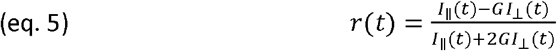

where *I*_‖_(*t*) and *I*_⊥_(*t*) are the parallel and perpendicular polarized fluorescence intensities, respectively, and *G* is the instrumental correction factor. For rotational diffusion in membranes, the anisotropy decay was fitted using a single exponential model

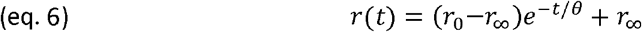

where *r*_0_ is the fundamental anisotropy, *r*_∞_ is the residual anisotropy reflecting restricted rotational mobility, and *θ* is the rotational correlation time^66,67^. The rotational diffusion coefficient was then obtained from

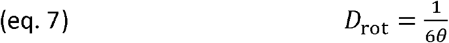

and used as an independent reporter of membrane viscosity.

Fluorescence correlation spectroscopy (FCS) was performed on the same setup as for FLIM/anisotropy as recently reported^41^. Cationic SUVs made of DOTAP:DOPE (50:50 mol ratio) were labelled with 0.1 mol% DOPE-Atto 655. Anionic SUVs made of DOPC:DOPS:Chol:PEG5000 (50-X:20:30:X), where X is the mol% of PEG5000, were labelled with 0.1 mol% Bodipy C_12_. Before each experiment, the collar ring of the water-immersion Olympus UPlansSApo (60x, 1.2 NA) objective was aligned by optimizing the molecular brightness of a solution containing Alexa488 and Alexa-647 fluorophores. These probes were used as a reference to calculate the confocal volumes for FCS, using 400 μm^2^/s as the values for free-diffusion in water^68^. To remove the contribution of detector afterpulsing, we corrected/removed background as background and fluorescence have different timing behavior in the TSCPS window. Fluorescence correlation spectroscopy (FCS) autocorrelation curves were fitted using a single-component three-dimensional diffusion model including a triplet-state contribution, given by

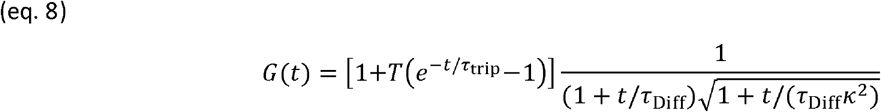

where *T* is the triplet-state fraction, *τ*_trip_ is the triplet lifetime, *τ*_Diff_ is the characteristic diffusion time, and *κ* is the axial-to-lateral radius ratio of the confocal volume (∼ 10). From the fitted *τ*_Diff_, the diffusion coefficient was calculated as

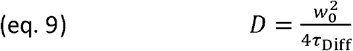

where *w*_*o*_ is the calibrated lateral focal radius. To estimate particle size, the hydrodynamic radius *r* was calculated using the Stokes–Einstein relation

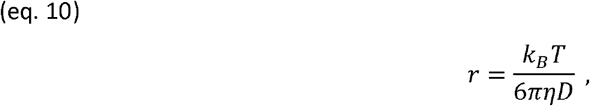

where *k*_*B*_ is the Boltzmann constant, *T* the absolute temperature, and *η* the viscosity of water (0.89 mPa·s).

Single-liposome fusion to supported lipid bilayers (SLBs) was imaged using a prism-type total internal reflection fluorescence (TIRF) microscope. In brief, DOPE-Rhodamine–labeled liposomes were excited using a 532 nm diode laser, while DOPE-Atto655–labeled membranes were imaged using a 640 nm solid-state laser. The evanescent field selectively illuminated liposomes within ∼100–200 nm of the SLB surface, enabling sensitive detection of docking, hemifusion, and full-fusion events at the single-particle level. Fluorescence signals were collected through a 60× water-immersion objective (UPlanSApo, Olympus) mounted on an inverted microscope (IX71, Olympus) and recorded using an electron-multiplying CCD camera (iXon 897, Andor Technology).

Fusion events were analyzed in two ways: First, to measure both interacting and weak or non-interacting behaviour, we tracked movies using the TrackMate tool in Fiji and custom analysis software on the stored XY-T traces as in Fig 2I. Second, for a more detailed analysis of the fusion events themselves, we developed custom analysis software in Python as follows: A typical fusion event is recognisable as an sharp increase in spot brightness, followed by a quickly expanding bright patch on the surface, followed by a full fading out of the spot or by disappearance of the particle. To quantify, we first detect event XY locations from a simple standard-deviation projection of the whole movie. Occurrences of events are typically low enough to not occupy the same XY on different time points. Around each position, we crop a XYT stack encompassing the event. From these, we use XT and YT projections of summed intensity to produce kymographs, that allows quick evaluation of the event as being a single release, a double release, a touch-and-go event and so on (see also Fig S8). Next, per time point we sample ring areas in XY from which to obtain summed intensity, similar to Ref^69^. Ring sizes are calculated such that each covers the same area as the central disk (with radius R0). For each larger ring, the summed intensity peaks slightly later in time, thus showing the passing diffusion front. The peak time of each ring relative to the event start should then scale linearly with the ring surface, with a slope reflecting D where D is the associated diffusion constant.

## Acknowledgements

The authors would like to thank Klaus Suhling for kindly donating the molecular rotor Bodipy C_12_; Anders Bart for support with acquisition and analysis of the spectroscopy data; and Dirk Bruin for his help with preliminary GUV experiments. We acknowledge funding from the EVOLF project (SUMMIT.1.004) which is financed by the Dutch Research Council (NWO).

## Author contribution

RBL designed the study, performed experiments, analysed data and wrote the manuscript. JK developed the pipeline for single liposome analysis. CD contributed lab infrastructure and edited the manuscript.

